# A new family of statistical tests for neuronal spiking and autocorrelated timeseries data

**DOI:** 10.1101/2023.10.30.564780

**Authors:** Jorrit S. Montijn, Guido T. Meijer, J. Alexander Heimel

## Abstract

Quantifying whether and when signals are modulated by autonomous or external events is ubiquitous in the field of neuroscience. Existing statistical approaches, however, are not ideally suited to do this, especially when the signals under scrutiny show temporal autocorrelations. For example, a standard approach in the analysis of calcium imaging data is to use a t-test on predetermined time-windows to quantify whether neurons respond (differently) to an event of interest. While this is attractive because of its simplicity, only average signal differences can be detected. In practice, neurons often show complex response dynamics which are missed by conventional statistical tests. More advanced methods, such as bin-wise ANOVAs, do not share this drawback, but can suffer from high false-positive rates or “ghost correlations” when applied to temporally autocorrelated data. To solve this issue, we have developed three novel statistical tests extending the original ZETA-test to other use cases: 1) a test for time-series data; 2) a two-sample test to detect differences in neural responses between two conditions; and 3) a two-sample test to detect differences in time-series data between two conditions. In addition, we have improved upon the original ZETA-test. We show that our methods have a statistical sensitivity superior to t-tests and ANOVAs and work well with temporally autocorrelated data where other approaches fail. Our methods are widely applicable and we present example applications to Neuropixels data, two-photon GCaMP imaging data, and human electrocorticogram data. Open-source code for implementations in MATLAB and Python is available on GitHub and PyPi.

**Significance Statement:** Neurophysiology involves the detection of weak signals in a noisy background. Often these signals, as well as the background, are temporally autocorrelated. This means that many statistical methods, such as the ANOVA, are inappropriate. In some cases, these statistical tests produce “ghost” signals that mislead researchers into thinking there is an effect of their experiment, when in reality there is none. We have therefore developed a new family of statistical tests, the ZETA-tests, that are not negatively impacted by temporal autocorrelations. They do not use arbitrary (hyper)parameters, like bin size, and show superior statistical sensitivity compared to other methods like t-tests and ANOVAs. We provide easy-to-use implementations and expect our methods to be useful for many neuroscientists from various disciplines.

## Introduction

Neurophysiological studies depend on a reliable quantification of whether and when neural signals are modulated by autonomous or external events, such as the presentation of a sensory stimulus. Early neurophysiology studies investigated response properties of a handful of single cells and depended heavily manual curation (Hubel & Wiesel, 1959; Mountcastle, 1957). Since these early days, the number of simultaneously recorded neurons has greatly increased together with statistical methods to analyse their activity, however, commonly used analytical methods are still lacking in various ways (Mesa et al., 2021). A common approach in neuroscience is to select a predetermined time-window and apply a statistical test to the mean level of activity in this period (Mazurek et al., 2014; Ringach et al., 2002). While such approaches are attractive because of their simplicity, statistical tests, such as the t-test, only detect differences in signal averages between the time windows of interest. On the other hand, more advanced model-based methods are rarely used, as they require fitting and manual hyperparameter tuning (Kass et al., 2014). These advanced methods can be very computationally and/or labour intensive, and may not be feasible for large data sets.

A middle ground between these approaches is perhaps the bin-wise ANOVA, when used in combination with an automatic bin-width optimization algorithm (Palm et al., 1988; Shimazaki & Shinomoto, 2007). While the ANOVA is an improvement over the t-test in many cases, it is ill suited to the analysis of temporally auto-correlated signals, as we will show later. In fact, most statistical methods assume some sort of sample independence; an assumption often violated in neural data. Such methods can therefore produce high false positive rates and “nonsense correlations” if applied to temporally auto-correlated signals that are otherwise unrelated to other signals (Harris, 2021; Meijer, 2021). An exception to this is a method commonly used in human electrophysiology, which uses a permutation-based clustering procedure (Maris & Oostenveld, 2007). This method is statistically sensitive and can be used on temporally-dependent data, but can only compare two sets of time-series data. Moreover, this approach still requires the selection of an arbitrary clustering threshold.

Neuroscience is therefore still lacking a general statistical test that requires no arbitrary parameters, binning, or manual curation, and is not negatively affected by temporally auto-correlated signals. Recent work has proposed such a statistical test for electrophysiological spiking data: the ZETA-test (Montijn et al., 2021). Several studies have already been published that use this approach. It is particularly well suited to detecting whether cells are driven by optogenetic stimulation in opto-tagging experiments (Dudok et al., 2021; Schneider et al., 2023; Spyropoulos et al., 2023; Szadzinska et al., 2021). It has also been used to detect response onset latencies (Oude Lohuis et al., 2022), and to quantify somatosensory and visual stimulus responsiveness (Burnett et al., 2023; Montijn et al., 2023; Qin et al., 2023; Ziegler et al., 2023). While the ZETA-test works well for spiking data, an important shortcoming is that it can only be applied to point events, such as spike times. Another limitation of the ZETA-test is that it cannot be used to determine differences between two conditions.

To address these limitations, we improved the ZETA-test and developed three new statistical methods: 1) a ZETA-test that can be applied to time-series data, like data from calcium imaging experiments. 2) A two-sample ZETA-test to determine whether there exists a difference in neuronal spiking activity between two conditions. 3) A two-sample ZETA-test for time-series data. We have tested the performance of these methods on real and synthetic data, and found that it outperforms common approaches such as t-tests and ANOVAs. We expect that our procedures may be of interest to statisticians and theoreticians, but we have written this article specifically with experimental neuroscientists in mind, who can use our open-source code in MATLAB (https://github.com/JorritMontijn/zetatest) and Python (https://github.com/JorritMontijn/zetapy) to obtain higher yields and more reliable results from their neurophysiological data.

## Results

### The original ZETA-test and the addition of a data-stitching step

The Zenith of Event-based Time-locked Anomalies test (ZETA-test) has been described in detail previously (Montijn et al., 2021) and in the Methods section, but we will concisely summarize its procedure here. First, we align all spikes to stimulus onsets, i.e. we build a raster plot (fig. 1A). Pooling all spikes across trials, we obtain a single vector of spike times relative to stimulus onset, and calculate the cumulative distribution as a function of time (fig. 1B). The deviation of this curve from a linear baseline represents whether the neuron has a higher or lower spiking density than a non-modulated spiking rate (fig. 1C, blue curve). We compare this pattern to the likelihood of observing it by chance by running multiple randomized bootstraps. In each bootstrap iteration, we jitter all stimulus-onset times to generate a single null hypothesis sample (fig. 1C, grey curves). After scaling the experimentally observed curve to the variation in the null hypothesis distribution obtained from all bootstraps, we transform it into a p-value by using the direct quantile position, or by approximation with the Gumbel distribution. Low ZETA-test p-values indicate that the neuron’s firing pattern is statistically unlikely to be observed if the neuron is not modulated by the event of interest.

**Figure 1.**
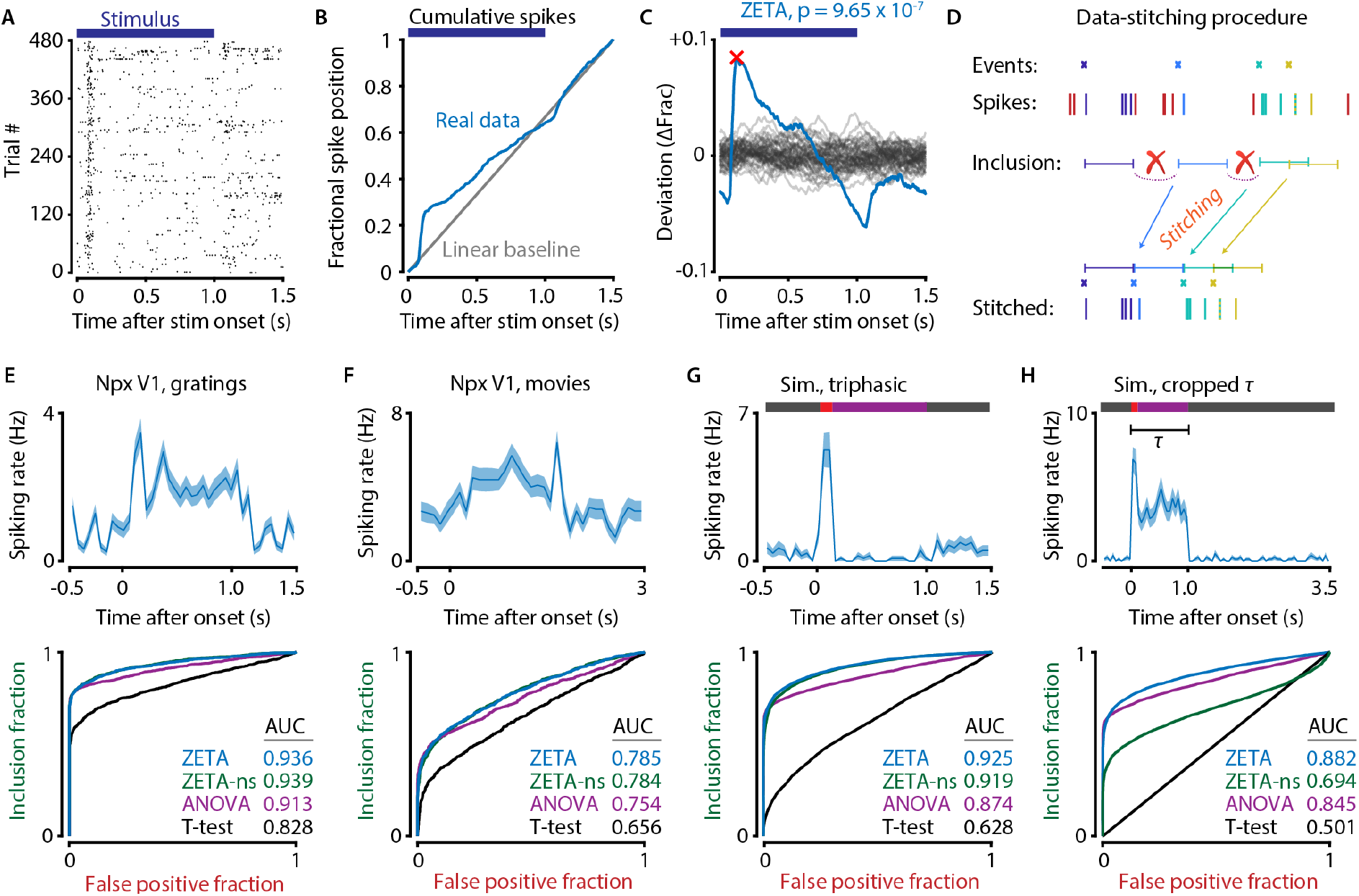
Data-stitching can improve the ZETA-test. A) Raster plot of an example neuron. B) Calculating the statistical metric for the ZETA-test depends on the difference between the real fractional spike positions (blue) and the null-hypothesis expectation from a constant, stimulus-unmodulated rate (grey). C) The Zenith of Event-based Time-locked Anomalies (ZETA, red cross) defines the significance after normalizing for the neuron’s intrinsic variability (grey curves). D) Overview of the new data-stitching procedure, where time periods are removed if they were not used for the calculation of the real deviation curve in panel B. E-H) Under specific conditions data stitching can dramatically improve ZETA’s performance. Top graphs show an example cell, bottom graphs show the statistical performance. ZETA indicates test with stitching, ZETA-ns without stitching. E,F) Experimental Neuropixels data of drifting gratings (E) and natural movies (F) shows the ZETA-test is superior to an optimal ANOVA and t-test, but stitching has little effect. G) Simulated neurons responding to stimuli with heterogeneous inter-trial intervals show the ZETA-test performs excellently, and the ZETA-test with data-stitching shows a small, but significant improvement. H) If the window of interest (τ) is chosen such that it discards transitions into and out of baseline activity, the sensitivity of the ZETA-test with no stitching is mediocre. Performing data-stitching improves the sensitivity of the ZETA-test to be superior to the ANOVA.

For all statistical benchmarking analyses hereafter, we will use a Receiver Operating Characteristic (ROC) analysis to quantify how well different statistical tests can discriminate whether a neuron is responsive to visual stimulation with drifting gratings. For the spike-based data sets, we used recordings with Neuropixels in the primary visual cortex (V1) of seven awake mice (Montijn et al., 2023). We analysed the responses of 1709 cells to drifting gratings and natural movies. In an ROC analysis, the true-positive rate (fraction of significantly responsive cells) is plotted on the y-axis, as a function of the false-positive rate (responsive cells after jittering the stimulus onset times), with each point corresponding to one p-value threshold. The ROC curve therefore follows the ratio of true-positives/false-positives for various values of α, ranging from the very lowest to highest p-value in the combined set of both true and false positives. For a properly calibrated statistical test, the false positive rate is equal to the significance threshold value α (commonly 0.05).

The first topic we investigated was a shortcoming in the original ZETA-test. The ZETA-test’s statistical model assumes that the real and null-hypothesis samples are taken from the same set of point events (spikes). While this holds true when stimulus events occur at fixed intervals, this will be false with heterogeneous inter-event durations, such as for self-generated behavioural events. In this latter case, jittering the onsets may lead to varying inclusion and exclusion of parts of the data (see Methods). We have therefore modified the ZETA-test to include an optional data-stitching step that ensures data conformity between the real and onset-jittered procedures (fig. 1D). While this change may seem significant from a purely theoretical point of view, it had surprisingly little effect on the statistical sensitivity in real-world experimental data with mild heterogeneity of inter-event durations (fig. 1E). We found the following statistical sensitivies: ZETA-test with stitching AUC = 0.936, ZETA-test without stitching AUC = 0.939, ANOVA AUC = 0.913, T-test AUC = 0.828. While the AUC of the ZETA-test was significantly higher than of the ANOVA (Mann-Whitney U-test, p = 1.4 × 10^−5^) and the t-test (p = 2.1 × 10^−74^), there was no difference in AUC between the stitched and non-stitched ZETA-tests (p = 0.53). For the ANOVA, we used the Shimazaki&Shinomoto procedure to calculate the optimal binning size (Shimazaki & Shinomoto, 2007), while for the t-test we tested compared the mean firing rate between the stimulus period (0.0 – 1.0 s) and pre-stimulus inter-trial interval (−0.5 – 0.0 s). We repeated the analysis on the responses to natural movies and found very similar results (fig. 1F). The ZETA-test outperformed the optimal ANOVA (p = 4.1 × 10^−4^) and t-test (p = 8.7 × 10^−45^), with no effect of stitching (p = 0.94).

Next, we tested the performance of these four tests under two different synthetic benchmarks with 10 000 simulated cells. In simulations for fig. 1G, we set the stimulus period to 1.0 s, the window of interest (τ) to 1.5 s, and varied the inter-trial intervals from 0.2 – 2.0 s to investigate the effect of the stitching procedure with heterogeneous inter-event durations. Under these conditions we found little effect of stitching: AUC = 0.925 vs 0.919 (p = 9.4 × 10^−4^) for with vs without stitching respectively. As before, the ZETA-test outperformed the ANOVA (p < 10^−100^) and t-test (p < 10^−100^). Secondly, we ran a simulation where we chose the window of interest τ such that it discarded the transitions into and out of baseline-level activity (fig. 1H). In this case, we found a large difference (p < 10^−100^) in statistical sensitivity when comparing stitching (AUC = 0.882) with no-stitching (AUC = 0.694). This result shows that stitching can be a critical step under some specific circumstances, especially for relatively sparse events where the window of interest may be unclear (for example, the duration of the period following licking).

### Time-series ZETA test

The ZETA-test in its original formulation can be applied to any two sets of point events, such as spike times and stimulus onsets. While its application to electrophysiological data is therefore straightforward, many other methods produce time-series data, such as calcium imaging. We previously applied the ZETA-test to transient-detected calcium imaging data and found that it performed quite well (Montijn et al., 2021). Nevertheless, a better and more generic solution would be a ZETA-test for time-series data, as this could also be applied to data obtained with EEG, patch recordings, and non-neurophysiological data. The Methods section describes an alternative formulation of such a time-series ZETA-test, which we applied to real data and various synthetic benchmarks to test its performance. In a nutshell, we replace the cumulative distribution of spikes by a cumulative sum of data values and perform some extra steps, such as data scaling (fig. 2A-E). An extra trick we employ is that if samples are acquired with varying latencies relative to the events of interest, this fact can be used to construct a super-resolution time-series average over trials. This allows the time-series ZETA-test to also effectively deal with data sampled with heterogeneous intervals.

**Figure 2.**
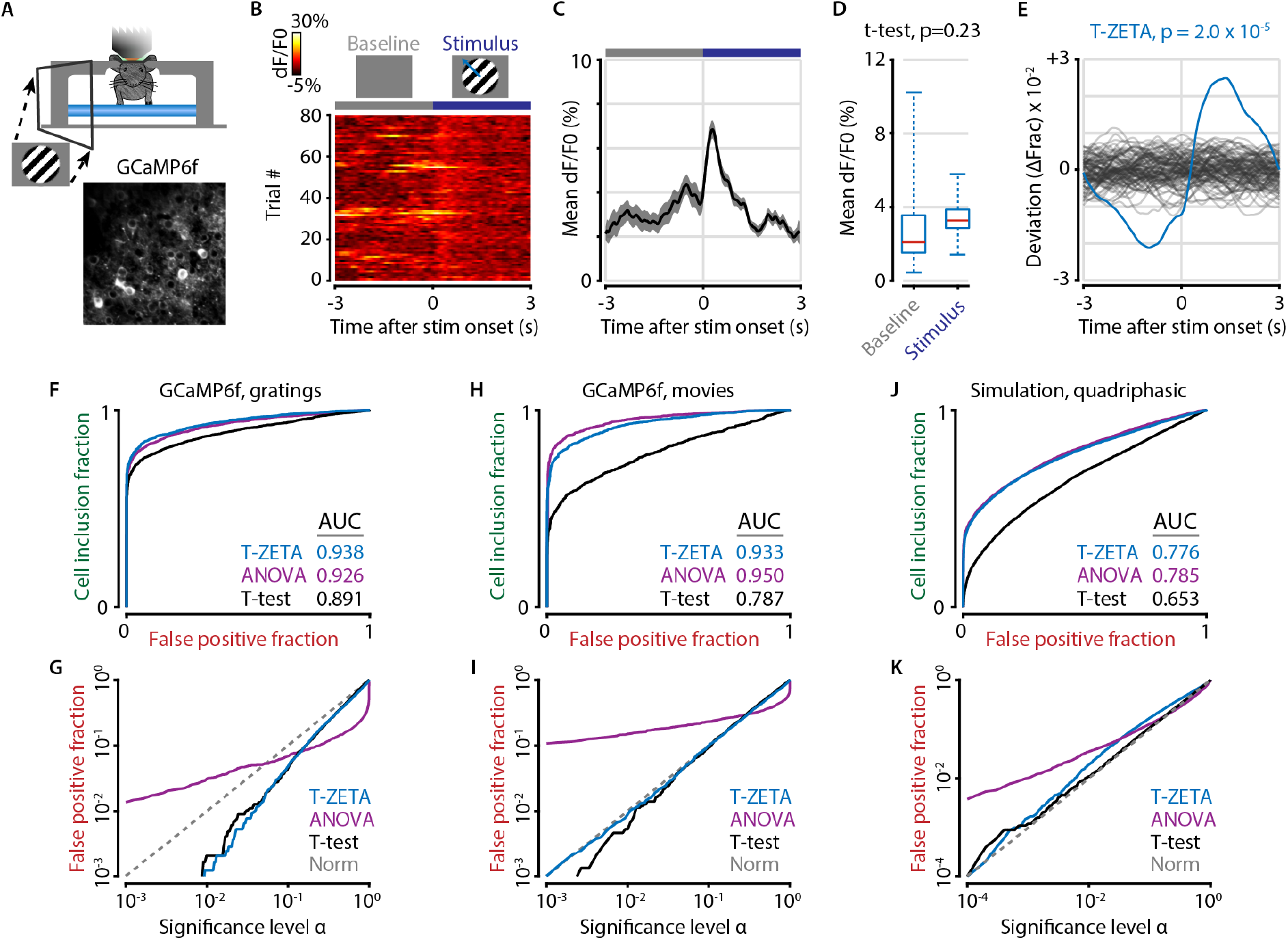
The time-series ZETA-test outperforms the t-test on statistical sensitivity, while the ANOVA applied to calcium data is excessively liberal and should not be used. A) The time-series ZETA-test can be applied to data recorded with calcium imaging. B) An example cell’s calcium activity (dF/F0) recorded with GCaMP6f in response to drifting gratings. C) This cell shows a clear onset response to the stimulus (period indicated by the blue bar), but also displays variable spontaneous activity and a lack of a sustained response. D) These factors reduce the difference in mean dF/F0 between the 3 s pre-stimulus baseline and 3 s stimulus periods, leading a t-test to erroneously classify this cell as non-responsive (paired t-test, n = 80 trials, p=0.23). E) Our alternative method (T-ZETA) does not use window-averages and that detects any time-locked deviations in neural activity. The blue curve shows the true deviation from a static level, and the grey curves show a subset of 50 bootstraps of deviations obtained by randomly jittering the stimulus onsets. F-K) ROC analyses to benchmark the statistical sensitivity under different conditions. F) Performance benchmark on GCaMP6f data of neurons responding to drifting gratings, showing the statistical sensitivity of the T-ZETA test (blue), the ANOVA (purple), and a t-test of 3s pre-vs 3s post-stimulus onset activity (black). G) Control analyses show the false positive rate (FPR) as a function of the threshold value alpha. The T-ZETA-test and t-test are somewhat conservative, as they lie below the theoretical norm (dotted line). On the other hand, the ANOVA is a poor statistical model when data points are not statistically independent: its p-value shows a highly non-linear relationship with the true probability that a sample is taken from the null-distribution and shows an excessively high false positive rate for α < 0.05. H,I) Same as F,G, but for natural movies, showing an even greater tendency of the ANOVA of produce excess false positives. J) We simulated quadri-phasic neurons with distinct baseline, onset, sustained, and offset responses where we filtered generated spike times with an exponential filter to mimic the effect of the slow dynamics of a calcium indicator. K) Simulations show one major cause of the ANOVA’s poor performance is a temporal correlated signal across bins. Here, the T-ZETA and t-test are close to the theoretical norm, but the ANOVA still shows an excess of false positives.

Like before, we used a receiver-operating characteristic (ROC) analysis to investigate the statistical performance of the time-series ZETA-test (T-ZETA). We tested the responses of 2430 cells to drifting gratings and natural movies. In the drifting grating data (fig. 2F), we found the T-ZETA-test outperformed the t-test (AUC = 0.938 vs 0.891, p = 1.0 × 10^−28^) as well as the ANOVA (AUC = 0.938 vs 0.926, p = 3.0 × 10^−3^). We further investigated the false positive rate of the three statistical tests, and compared this to the theoretical norm (fig. 2G). If a statistical test is well-calibrated, the number of false positives in a randomized data set should be roughly equal to the significance level α. The t-test and T-ZETA-test showed false positive rates lower than the theoretical norm, meaning they were somewhat conservative. While not ideal, this behaviour is acceptable, as it does not lead to false discoveries of an effect where there is none. On the other hand, the ANOVA showed a rather problematic false-alarm rate; it was too liberal at low p-values and too conservative at higher ones (fig. 2G). For example, to obtain an empirical false positive rate of 0.001 using the ANOVA, one would require a p-value threshold α of around 10^−13^ rather than the expected 10^−3^ if p-value scaling had followed the theoretical norm. In the natural movie data, we found similar results (fig. 2H). Here, the T-ZETA-test outperformed the t-test by a considerable margin (AUC T-ZETA = 0.933, t-test = 0.787; p < 10^−100^). Although the ANOVA’s AUC (0.950) was higher than the T-ZETA’s (p = 3.7 × 10^−7^), the ANOVA again showed a strong excess of false positives (fig. 2I), rendering the ANOVA unsuited for the application to this data set.

We suspected the excess false positive rate of the ANOVA is caused by the ANOVA’s model assuming statistical independence between adjacent bins of the peri-stimulus time histograms (PSTHs). As calcium indicators are low-pass filters, and even the underlying spiking rate itself is not temporally independent, this assumption is violated. To confirm that a simple temporally auto-correlated signal leads to excess false positives in the ANOVA, we ran a simulation with 10 000 quadriphasic cells. We generated spiking times using exponential inter-spike intervals that we varied over the course of a single trial, and applied a Gaussian-filter on spike counts of 40-ms bins (25 Hz) to simulate the effect of the calcium indicator. The results of this simulation were similar to those we found in real data: the AUC of the T-ZETA and ANOVA were similar (fig. 2H), but the false-positive rate of the ANOVA was very high (fig. 2K). We therefore strongly advise against using the ANOVA, or other bin-based analyses, when analyzing temporally autocorrelated time-series data.

### Two-sample ZETA test

We developed the ZETA-test to detect whether a neuron’s activity was modulated by the occurrence of a set of events. This approach works especially well to, for example, determine the genetic subtype of neurons in an opto-tagging data set (Dudok et al., 2021; Schneider et al., 2023; Szadzinska et al., 2021). Ideally, however, one would also be able to use a binless statistical test to determine whether two neurons behave differently in response to the same stimulus, or whether the same neuron differs in response between two stimuli. We have therefore developed a two-sample zeta-test which can be used to assess the difference in response between two conditions. In short, rather than comparing the response of a neuron to a linear baseline rate, we define a temporal spiking-density deviation vector as the difference in normalized cumulative spike rates between two conditions (fig. 3A-D). We compare the maximum absolute deviation to those we obtain using a trial-swapping procedure where we re-generate a deviation vector in every bootstrap-iteration in which we randomly assign trials to one or the other condition. The statistical significance of the test is then the likelihood to find the real data within the distribution of randomized bootstrapped controls.

**Figure 3.**
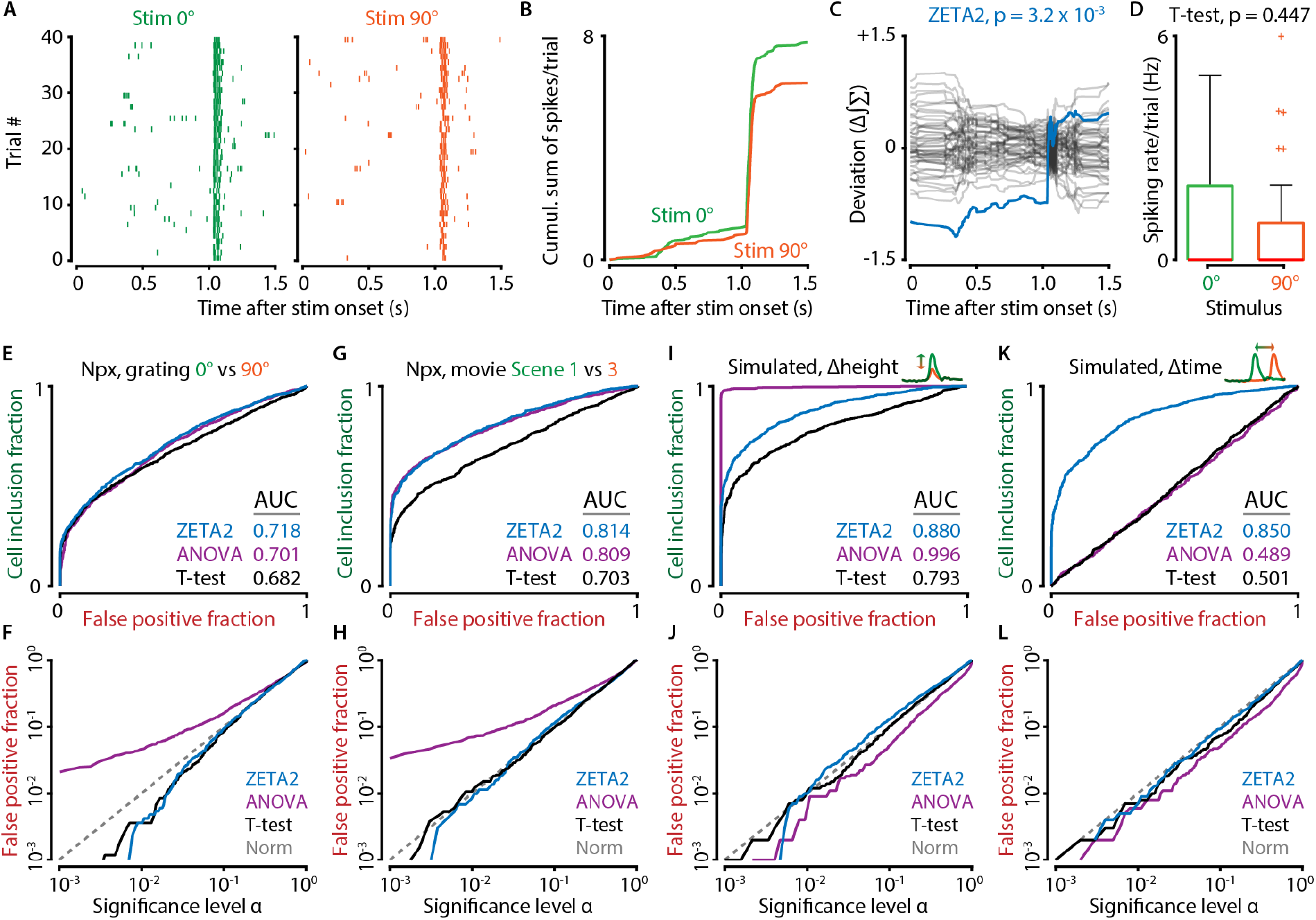
The two-sample ZETA-test quantifies response differences between two conditions. A) Example spike raster plots of one neuron responding to two different sets of drifting grating stimuli. B) The statistical metric for the two-sample ZETA-test depends on the difference in the cumulative sum of the spikes per trial for the two conditions. C) The real difference (blue) is compared to the difference obtained using resamplings where trials are randomly assigned to condition 1 (0 degree direction) or 2 (90 degree direction). D) The two-sample ZETA-test (ZETA2) detects a difference between these two responses (p = 2.7 × 10^−6^), but a two-sample t-test does not (p=0.158). E-L) Various benchmarks to compare the performance of the ZETA2-test to an optimally-binned two-way ANOVA and t-test. E) Comparison of the response of all 1504 neurons in V1 recorded with Neuropixels to drifting gratings moving in the 0 degree or 90 degree direction. The ZETA2-test shows a similar statistical sensitivity to the ANOVA (p = 0.071) and higher than the t-test (p = 1.7 × 10^−4^). F) The fraction of false positives in a shuffle-control is close to the theoretical norm for the ZETA2-test and t-test, but the ANOVA again shows an excessively high false positive rate. G,H) As E,F, but for natural movie scene discrimination. G) The ZETA2-test shows a significantly higher statistical sensitivity than the t-test (p = 5.1 × 10^−30^), but is not different from the ANOVA (p=0.52). H) The ANOVA shows an excess false positive rate. I-L) Simulation with exponential-ISI neurons of the best-case (I,J) and worst-case (K,L) scenario for the ANOVA, where the difference between two conditions is defined only by the number of spikes in a short response peak (I,J) or only by a 2 ms difference in peak time (K,L). The ANOVA performs excellently in its best-case scenario, but performs at chance-level for the worst-case scenario. The ZETA2-test, in contrast, performs well in both cases. Note that in these simulations, in the absence of temporally correlated spiking, the ANOVA’s false positive rate is close to the theoretical norm.

We tested the performance of the two-sample ZETA-test (ZETA2-test) using multiple benchmarks. First, we used data from Neuropixels recordings in primary visual cortex (Montijn et al., 2023). We ran two comparisons on these data: first, we tested whether a neuron’s response differed between two sets of stimuli (gratings drifting at 0 and 90 degrees) (fig. 3E). We compared the performance of the two-sample zeta-test to that of the two-sample t-test, and a two-way ANOVA. For the ANOVA, we took the p-value to be the lower of the two Bonferroni-corrected p-values for the main effect between conditions (i.e., whether there was a mean-rate difference) and the *bin x condition* interaction effect. We determined the optimal bin size for the ANOVA using the Shimazaki & Shinomoto procedure after pooling spikes from both conditions (Shimazaki & Shinomoto, 2007). The ZETA2-test performed best (AUC = 0.718), followed closely by the ANOVA (AUC = 0.701; p = 0.071), and less closely by the t-test (AUC = 0.682; p = 1.7 × 10^−4^). We investigated the false positive rates of these three tests and found that the ANOVA again showed an excess of false positives (fig. 3F). Secondly, we quantified the performance of these methods on testing the difference in response to natural movie scenes (scene 1 vs scene 3) (fig. 3G). The AUCs of the ZETA2-test and ANOVA were similar (0.814 and 0.809 respectively; p = 0.52), followed at some distance by the t-test (0.703; p = 5.1 × 10^−30^). Similar to the drifting grating data, the ANOVA showed a high false positive rate, while the ZETA2-test and t-test were close to the theoretical norm (fig. 3H).

Next, we simulated two cases to highlight the strong and weak points of the ZETA2-test. We generated 1000 pairs of cells with a randomly assigned background spiking activity level (mean = 1.0 Hz) following an exponential inter-spike interval distribution. Both cells always had the same average background spiking rate, but on top of these spikes we added a single spike in 25% of 240 trials for condition 1 and 50% for condition 2 (with a temporal delay of exactly 55 ms and a trial-to-trial jitter of 1 ms). The only difference between these cells is therefore the spike count during the response peak. The optimal ANOVA could select a bin width that maximized the spike-count differences and yielded an impressive AUC of 0.996 (fig. 3I). The ZETA2-test followed with 0.880 and the t-test with 0.793. As expected from simulated cells that show no temporally auto-correlated spiking behaviour, the false positive rate of the ANOVA was close to the theoretical norm (fig. 3J). Finally, we simulated a similar situation where there was no difference in peak height, but only in peak time. For condition 1 we set the peak time at 53 ms and for condition 2 at 55 ms, adding a single spike in 25% of all trials. The ZETA2-test was well able to differentiate these cells with an AUC of 0.850 (fig. 3K,L). The optimal ANOVA was near chance with an AUC of 0.489, as the selected bin size was often too large, and the t-test had an AUC of 0.501. This shows that the two-sample ZETA test excels at detecting whether neurons show different temporal modulations of their firing rates between conditions.

### Two-sample time-series ZETA test

The fourth member of the ZETA-family of tests is the two-sample version of the time-series ZETA-test (T-ZETA2-test). As for the spike-based ZETA2-test, we compared the performance between a two-way ANOVA, two-sample t-test and T-ZETA2-test. Moreover, we also tested a statistical clustering-based method popular in the field of human electrophysiology (Maris & Oostenveld, 2007). We adapted this Maris & Oostenveld cluster permutation test, which we will abbreviate to the “Clust-t” test, to obtain a p-value for the difference in response between two sets of time-series data, and compared its performance to those of the other tests (fig. 4).

**Figure 4.**
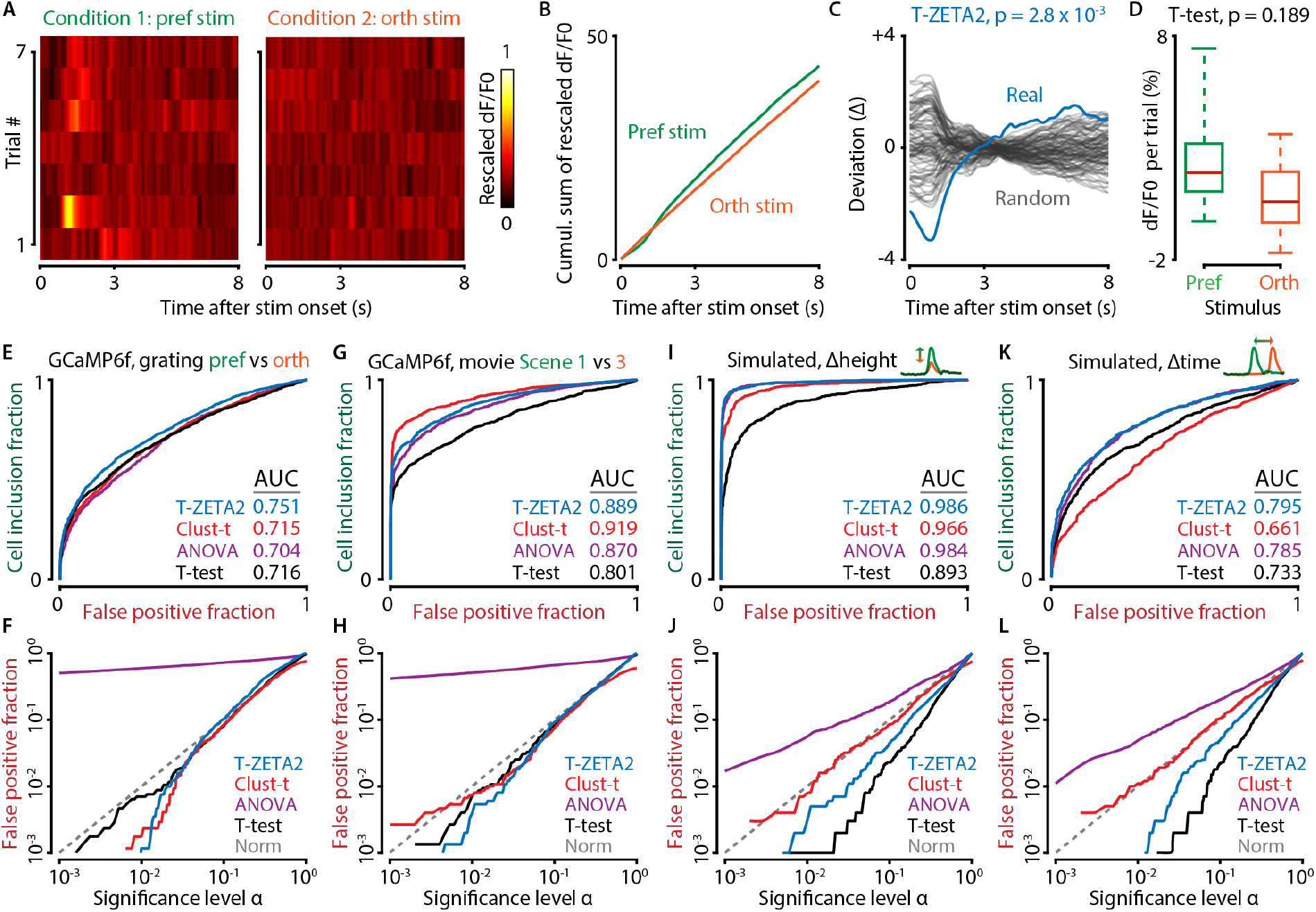
The two-sample time-series ZETA-test (T-ZETA2) can discriminate neural activity in calcium imaging data. A) Heat maps of one example V1 neuron’s response to drifting gratings moving in its preferred and orthogonal direction. B) The T-ZETA2-test uses the cumulative sum of rescaled neural activity (here dF/F0). C) The deviation in cumulative sum defines the ZETA-metric and is compared to deviation curves obtained after randomly combining trials from either condition, similar to the spike-based ZETA2-test. D) In this example, the T-ZETA2-test detected a significant difference in responses between the two conditions (p=2.8 × 10^−3^) but a t-test did not (p=0.189). E) Benchmark discriminating between the responses of two neurons to the same set of stimuli, including the “Clust-t” Maris & Oostenveld test, a popular method in human electrophysiological studies. The T-ZETA2 shows a higher AUC than all other tests: T-ZETA2 vs t-test, p = 4.3 × 10^−5^; vs ANOVA, p = 4.2 × 10^−8^; vs Clust-t, p = 2.4 × 10^−5^. F) A shuffle-control analysis of the false positive rate reveals that the ANOVA shows an excessively high false positive rate, rendering the p-values it returns practically meaningless. For example, at an α of 0.05 the ANOVA has a 68% false positive rate (rather than 5%). The T-ZETA2-test, Clust-t test, and t-test show false positive rates close to the theoretical norm. G,H) Same as E,F, but applied to neuronal responses to natural movie scene transitions. Here, the Clust-t test performs best and the t-test performs worst, while the ANOVA still shows excessively high false positive rates. I-L) Simulations of peak-response difference (I,J) and peak-time difference (K,L) reveal the T-ZETA2-test performs best on these benchmarks, with the Clust-t test performing rather poorly. Across different conditions, the T-ZETA2-test and Clust-t test show the highest AUCs and well behaved false-positive rates, but which of these two is more appropriate depends on the specifics of the underlying data set.

We benchmarked all four tests by applying them to the same calcium imaging data sets as for the one-sample time-series ZETA-test, now calculating whether there is a difference in the response of one neuron to its preferred drifting grating direction and the stimulus orthogonal to it (fig. 4E,F). To compute the baseline false-positive rate, we split the preferred stimulus trials into two random sets of trials and calculated whether there was a difference in response between them. We found that the AUC was highest for the T-ZETA2-test, followed by the t-test, Clust-t test, and ANOVA (AUCs of 0.751, 0.716, 0.715, and 0.704 respectively). The performance of the T-ZETA2-test was significantly higher than the three other tests (p = 4.3 × 10^−5^, p = 2.4 × 10^−5^, p = 4.2 × 10^−8^), while the Clust-t test did not differ in performance from the t-test and ANOVA (p = 0.89, p = 0.22). We also tested the performance on natural movie responses (Scene 1 vs Scene 3), but found that in this case the Clust-t test performed best, followed by the T-ZETA2, ANOVA, and t-test (AUCs of 0.919, 0.889, 0.870, and 0.801 respectively; fig. 4G,H). All pairwise performance comparisons were significant (p < 2.0 × 10^−3^). In both cases, the false positive rate (FPR) for the ANOVA was again exceptionally high: at an α of 0.05 the FPR was 0.677 for the grating comparison and 0.608 for the natural movie comparison. In contrast, the three other tests had false alarm rates close to the theoretical norm. As we noted above, and confirmed here, the ANOVA is very clearly the wrong statistical model for the analysis of time-series data due to the temporal autocorrelation of signals across bins.

We further investigated the behaviour of the four methods using the synthetic benchmarks we described in the previous paragraph: we simulated cells with either a difference in peak height (fig. 4 I,J), or in peak time between two conditions (fig. 4 K,L). As for the one-sample time-series data, we applied an exponential filter to spike counts in 40-ms bins (25 Hz). In the Δheight simulation, the T-ZETA2-test performed best, followed by the ANOVA, Clust-t, and t-test (AUCs of 0.986, 0.984, 0.966, and 0.893 respectively). Interestingly, the Clust-t test performed rather poorly in the Δtime simulation; the T-ZETA-2-test performed best, followed by the ANOVA, t-test, and at some distance the Clust-t test (AUCs of 0.795, 0.785, 0.733, and 0.661 respectively). The ANOVA showed a high false positive rate in both simulations, but to a lesser degree than in the experimental data sets.

Overall, the performance of the two-sample time-series ZETA-test was good and robust: it showed a false-positive rate close to the theoretical norm and a high statistical sensitivity in all four cases we investigated. The Clust-t test also performed well, especially in the natural movie data set, but it appears more sensitive to specific properties of the dataset it is applied to. While the T-ZETA2-test performed best in three of four cases and second-best in the natural movie data set, the Clust-t test performance was more variable. It performed excellently in the natural movie data set, but was surpassed by even the t-test in the Δtime simulation. Moreover, the ANOVA’s performance in terms of AUC was reasonable, but its exceptionally high false positive rate renders that moot; with a false positive rate of 0.68 at α = 0.05, it may actively mislead researchers to think they found an effect where there is none.

### The T-ZETA2-test localizes the fusiform face area in human ECoG data

The potential applications of the ZETA-family of statistical tests go beyond cellular neurophysiological data. The time-series ZETA-tests can theoretically be applied to any kind of temporally evolving data, neuroscientific or otherwise. We developed the ZETA-tests mainly with cellular neuroscience in mind, but the time-series ZETA-test may also be applied to, for example, human electrocorticogram data (ECoG). We analysed an open data set, where participants were shown pictures of houses and faces while ECoG signals were recorded from various brain locations (Miller et al., 2016). These locations varied between patients and depended on the estimated focal loci of their epileptic seizures. The questions we asked were: 1) how many responsive electrode sites does the T-ZETA-test detect, and does this exceed other approaches? 2) can the T-ZETA2-test be used to detect which sites are differentially responsive to house and face stimuli?

We applied three approaches to detect whether a recording site responded to house stimuli: the t-test, T-ZETA-test, and a procedure described by the original authors, which uses a linear model to estimate the amount of variability in the neural signals that can be explained by the stimuli (R^2^) (Miller et al., 2016). We found that the T-ZETA detected a response in 78% of electrode sites across all subjects, which exceeded the t-test at 58% (p = 2.6 × 10^−4^) and the R^2^ procedure at 39% (p = 9.3 × 10^−13^) (fig. 5A-C). False alarm rates were around 5% for the T-ZETA and t-test (6.8% and 4.7% respectively), but the R2 procedure appeared overly conservative with an FPR of 0%. Moreover, as expected, occipital recording sites were responsive to these visual stimuli (fig. 5D). We repeated the same analysis for face stimuli, and found similar results (fig. 5E-G). The T-ZETA detected a response in 80% of electrode sites, which again exceeded the t-test at 44% (p = 4.0 × 10^−12^) and the R^2^ procedure at 45% (p = 1.6 × 10^−13^), and FPRs were similar to before (4.6%, 5.1% and 0% respectively). The pattern of responsive sites for face stimuli was also similar to that for house stimuli, showing the strongest responses in occipital regions (fig. 5H).

**Figure 5.**
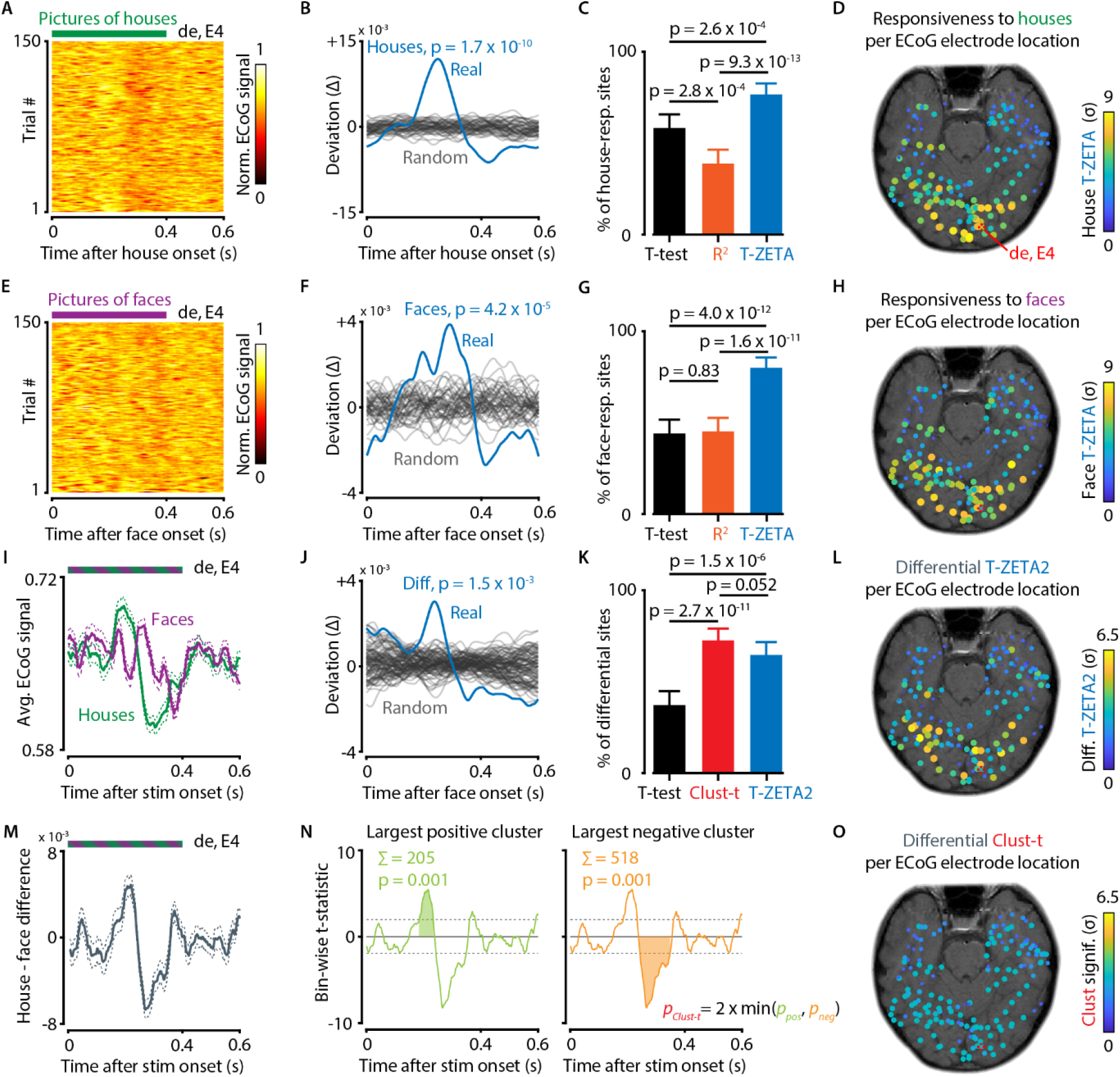
The time-series ZETA-tests can be applied to a variety of data types, including human electro-corticogram (ECoG) data. A) Heat maps showing the normalized ECoG signal in response to a picture of a house for one example electrode. B) The T-ZETA-test can be used to test whether an electrode is responsive to house stimuli; this example electrode showed a highly significant response (p = 1.7 × 10^−10^). C) Applying a t-test, signal predictability procedure (R^2^), and T-ZETA-test, we found that the T-ZETA-test was the most statistically sensitive, detecting a significant response to house stimuli in 78% of electrode sites across all subjects. The t-test detected a responses in 58% of electrodes, followed by the R^2^ procedure with 39%. D) Map showing approximate electrode locations. Colour indicates the T-ZETA’s statistical significance of the electrode’s response. The example electrode is marked in red. E) Example electrode’s response to different pictures of faces. F) T-ZETA-test applied to this electrode’s response to face stimuli (p = 4.2 × 10^−5^). G) As C, but for face stimuli. Site responsiveness: T-ZETA-test, 80%; t-test, 44%; R^2^, 45%. H) T-ZETA responsiveness map for face stimuli. I) Mean +/-SEM of the signal averaged over stimulus presentations for house stimuli (green) and face stimuli (purple). J) While this electrode is responsive to both types of stimuli, it also shows a significant difference in response between the two stimulus types (T-ZETA2-test, p = 1.5 × 10^−3^). K) Testing whether an electrode showed a different response to houses and faces, we found the following differential responsiveness percentages: T-ZETA2-test, 63%; t-test, 37%; Clustering procedure, 73% (see M,N). L) T-ZETA2 differential responsiveness to houses vs. faces. Note that many occipital locations are less significant than in D,H, and the strongest responses match the approximate location of the fusiform face area. M) The average difference in response between house and face stimuli. N) A clustering test often used in human electrophysiology uses the largest cluster-based sum of bin-wise t-statistic values (Maris & Oostenveld, 2007). This procedure also yielded a significant difference in responses between houses and faces (Clust-test, p = 2.0 × 10^−3^). O) Differential responsiveness of the clustering test to houses vs. faces. The maximum significance is limited by the number of bootstraps (here n = 1000), as the p-value is defined as the quantile position of the real data in the shuffled samples.

Earlier work has shown that temporal brain regions, such as the fusiform face area, respond specifically to faces (Sergent et al., 1992; Kanwisher et al., 1997). We therefore next investigated whether we could use the T-ZETA2-test to pinpoint the anatomical locations where these differences in neural responses to houses and faces are strongest (fig.5 I-O). We compared the differential responsiveness using the T-ZETA2, Clust-t, and t-test, and found T-ZETA2-test (63% differentially response) performed close to the Clust-t test (73%, p = 0.052), and better than the t-test (37%, p = 1.5 × 10^−6^). Interestingly, the anatomical locations where the T-ZETA2-test detected the strongest differential responsiveness indeed matched the approximate location of the fusiform face area (fig. 5L), while this pattern was less obvious using the Clust-t test (fig. 5O). These results show that the T-ZETA2-test and the Clust-t test have complementary qualities and comparable statistical performance. Which test is better therefore depends on the specifics of the underlying data, as well as the question under investigation.

## Discussion

We presented new members of the ZETA family of statistical tests that can be applied to two-sample comparisons and time-series data. The family of ZETA-tests is built upon a statistical method for determining whether neuronal responses are modulated by the occurrence of events. The method is sensitive to complex response patterns and robust in the face of temporally autocorrelated signals – a situation where the ANOVA shows an excessively high false alarm rate (fig. 6A). In all experimental data, the ZETA-tests showed markedly improved statistical sensitivity and/or robustness compared to established and powerful statistical techniques, such as t-tests and optimally-binned ANOVAs (fig. 6A-E). We also compared the T-ZETA2-test with the Maris & Oostenveld clustering-based method, which is often used in human electrophysiology. While statistical sensitivity was similar and the specifics of the underlying data determine which test performs better (fig. 4,5), the T-ZETA2-test has three distinct advantages: (1) it does not require an arbitrary cluster threshold (fig. 6F-I), (2) outputs unbounded p-values, and (3) requires fewer bootstraps iterations. In conclusion, the family of ZETA-tests is easier to use than established methods, as it can be applied directly to raw spike times or time-series data and stimulus onsets, and the lack of parameter selection naturally lends itself to the bulk-analysis of large numbers of cells.

**Figure 6.**
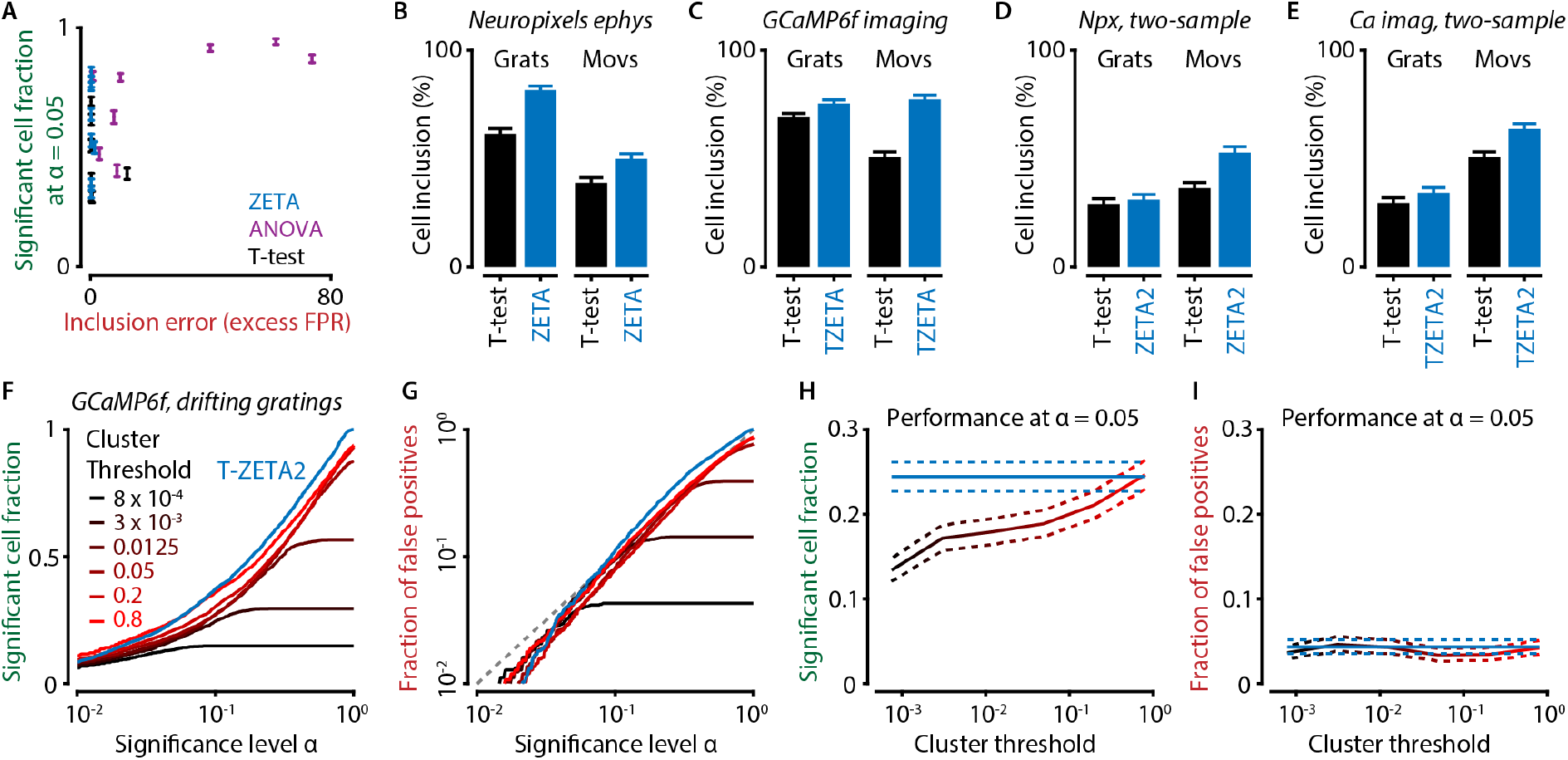
Members of the ZETA-family of statistical tests perform well on different types of data and outperform common statistical methods. A) Summary of the performance of the ZETA-family of statistical tests, ANOVAs, and t-tests. The ZETA-family and t-test show well-calibrated behaviour close to the theoretical norm, while the assumption of bin-wise independence of the ANOVA leads to excessively high false positive rates. B-E) Of the two types of tests that show well-calibrated false positive rates, the ZETA-family tests exceed the t-test’s inclusion rate at α = 0.05 in each experimental data set we investigated. Grats: drifting grating stimuli. Movs: natural movie stimuli. F-I) The Maris & Oostenveld clustering test performs well on ECoG data, but shows variable performance on other types of data. F) The clustering test requires the selection of a clustering threshold, an arbitrary hyperparameter. Running the same benchmark as in Fig. 4E,F, we found the T-ZETA2-test outperformed the clustering test for all except for the highest cut-off (0.8), where performance was equivalent. G) False positive rates of all tests were close to the theoretical norm, but note that low cluster thresholds can lead to the absence of significant clusters, causing overly conservative behaviour at high α values. H) The T-ZETA2-test showed a higher fraction of significant cells at an α of 0.05 than the Maris & Oostenveld clustering test for all cluster thresholds, except 0.8. Note that the default is threshold is 0.05, and that these results are not corrected for multiple comparisons. I) The false positive rate was close to 0.05 in all cases.

We described the procedure of data-stitching to deal with heterogeneous inter-event durations (fig. 1). While this procedure is in most cases as good as, or better than, using the unstitched data, there are exceptions. In the situation where events with heterogeneous inter-event durations lead to short responses in an otherwise stationary background activity, data stitching may reduce the statistical sensitivity of the ZETA-test. Extending the jittered data into periods of stationary inter-event epochs means the variance of the jittered data is reduced, compared to stitched data where larger jitter magnitudes can include the response of the preceding or following event. However, when a set of control data exists where the only difference from the experimental data is the presence of the events of interest, it might be preferable to instead use the two-sample ZETA-test and directly compare these two conditions with surrogate events for the control condition.

The T-ZETA-test described in this paper may superficially resemble the Kolmogorov-Smirnov (KS) test, so one could wonder how these approaches differ. In our original derivation of ZETA, we perform an in-depth comparison between the KS-test and ZETA-test, so we refer to the reader to this earlier work for more details (Montijn et al., 2021). In short, the main difference between ZETA and KS is that the KS-test is very sensitive to any difference in the cumulative distribution between two conditions, even if that difference is not time-locked to a stimulus, but results from intrinsic differences in the distribution of neuronal activity. This makes the KS-test unsuitable for application to neuronal activity, as it generates many false positives. The ZETA method takes only the maximum deviation as a metric for “differentness”, and as a result becomes less sensitive to the exact shape of the cumulative distribution.

The ZETA2 test is also much more sensitive than the two-sample Kolmogorov-Smirnov test for data with multiple trials, as the Kolmogorov-Smirnov test does not consider the variability across trials. The ZETA2 test also shows some similarities to a standard permutation test over binned data. Here, the main difference is that each null hypothesis sample in the case of the ZETA2 test is the maximum deviation over all time points rather than a value for each sample time, permuted over trials. The ZETA2 test therefore circumvents the multiple-comparison problem that would arise if one would, for example, perform a permutation test for each sample time. Because of this invariance to the number of sample points, we were also able to use some other tricks, such as creating a super-resolution reference time vector to allow for its application to data with heterogeneous sampling intervals.

In conclusion, members of the ZETA family are simpler, more statistically powerful, and less error-prone tools than bin-based PSTHs, t-tests and ANOVAs. ANOVAs can be powerful tools when combined with optimal-binning algorithms, but assume statistical independence between bins. As our results have shown, in many cases this causes the ANOVA to output misleading p-values. The expanded utility of the ZETA-test to two-sample comparisons and time-series data can therefore provide an attractive, more robust, and more statistically sensitive alternative to other approaches. Implementations of the four ZETA-tests described in this paper are available in MATLAB (https://github.com/JorritMontijn/zetatest) and Python (https://github.com/JorritMontijn/zetapy).

## Materials and Methods

### A summary of the ZETA-test

The Zenith of Event-based Time-locked Anomalies (ZETA) test has been described in detail previously (Montijn et al., 2021), but for completeness we will describe it below. In the paragraphs thereafter, we will present a modification that addresses a potential shortcoming of the original ZETA-test, explain how we constructed a test for time-series data, and present statistical procedures to perform two-sample comparisons for both spike-based and time-series data.

The metric *ζ* is computed on a vector of *i = [1 … n]* spike times ***x***, and a vector of *k = [1 … q]* event times ***w***. First, we make a vector ***v*** of the spike times in ***x*** relative to the most recent stimulus onset, as when making a raster plot of spike times:

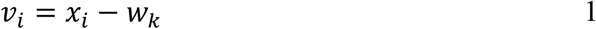

Where *k* is chosen such that

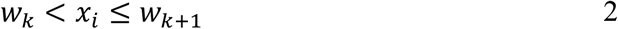

Next, we remove all spike times that are larger than a cut-off value *τ*, for example the trial-to-trial duration, and add two artificial spikes at t = 0 and t = *τ* to ensure the procedure is well defined in the case of one spike. We sort the *n* spike times in ***v*** such that *v*_*i*_ *< v*_*i+1*_, and calculate the fractional position *g*_*i*_, ranging from 1/*n* to 1, of each spike time in ***v***:

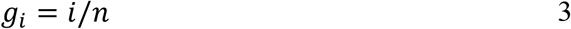

Therefore, ***g*** represents a neuron’s cumulative density function sampled at the spike times in ***v***. In order to quantify whether this distribution is different from our null hypothesis – i.e., that the neuron’s firing rate is not modulated with respect to the stimulus onset – we compare this vector to a linear baseline density vector ***b***. If a neuron’s spiking rate is constant, the cumulative density function is linear over time, and therefore the expected fractional position of spike *i* at time *v*_*i*_ converges to the spike time divided by the trial duration *τ* as the number of events *q* increases:

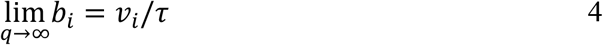

The difference *δ*_*i*_ between *g*_*i*_ and *b*_*i*_ therefore gives a neuron’s deviation from a temporally non-modulated spiking rate at time point *v*_*i*_:

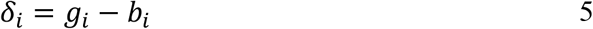

As shown previously, using *δ*_*i*_ to compute ZETA would make it dependent on the choice of onset times (Montijn et al., 2021). Therefore, we create ***d***, a time-invariant mean-normalized version of ***δ***:

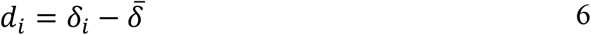

Where

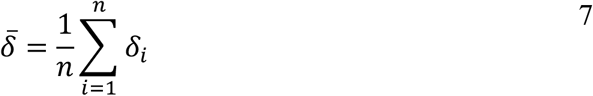

We then define the Zenith of Event-based Time-locked Anomalies (ZETA, or *ζ*_*r*_) as the most extreme value, i.e. the maximum of the absolute values:

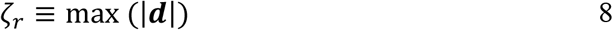

Having calculated ZETA from the temporal deviation vector ***d***, we wish to quantify its statistical significance. We therefore construct a null hypothesis distribution by repeating the above procedure *M* times with shifted event-times ***w’***, where for each jitter iteration *m* and event *k*, we move each event time by a random amount sampled from the interval [-τ, τ]:

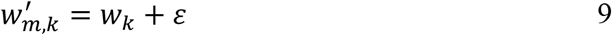

Where each *ε* is independently drawn from a uniform distribution *U*:

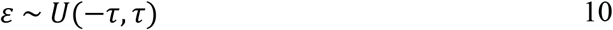

We repeat this process *M* times; where for each jitter iteration *m*, we calculate ***δ****’(m)*:

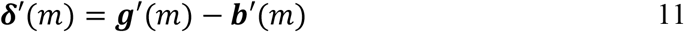

As before, we mean-normalize ***δ’****(m)* and define a null-hypothesis ZETA sample *ζ’(m)* as:

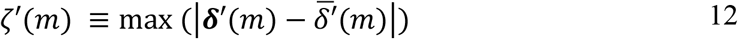

Having a way to generate null-hypothesis samples, we can now calculate the significance of *ζ*_*r*_ either directly, using the quantile position of *ζ*_*r*_ in the set of *M* samples ***ζ’***, or we can approximate it by using the Gumbel distribution estimated from these samples (Montijn et al., 2021). Either way, we obtain an estimate that asymptotically converges to a deterministic value as the number of jitter iterations *M* grows. We then use the standard normal’s quantile function Φ^−1^ to obtain a corrected ZETA metric ζ that is interpretable as a z-score:

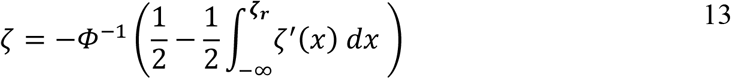

With its corresponding p-value defined by:

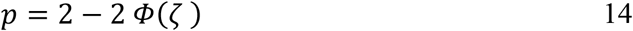

Where *Φ* is the cumulative normal distribution. Note that when we refer to ZETA or ζ in the rest of the manuscript, we mean the corrected version and its p-value as defined above.

### Data-stitching to ensure conformity between the real data and random bootstraps

One step that was not present in the original description and implementation of the ZETA-test, was to discard any data that was outside all event-window bounds (*w*_*k*_, *w*_*k*_ *+ τ*) for all events *k=1… q*. In the original definition, a spike time *x*_*i*_ for which *w*_*k*_ *+ τ < x*_*i*_ *< w*_*k+1*_ holds true, was not included in the calculation of the real zeta metric, but could be included in the random resamplings if also *w*_*k*_ *+ τ < x*_*i*_ *< w*_*k*_ *+ 2τ* or *w*_*k+1*_ *-τ < x*_*i*_ *< w*_*k+1*_. Note that this will only occur if *τ* is chosen to be shorter than the longest inter-event duration. In the original definition of ZETA, we implicitly assumed a fixed inter-event duration equal to *τ*, but this does not always hold true. For example, when applying the ZETA-test to self-generated behavioural events such as licking, inter-event durations will be highly variable.

Following the original ZETA definition, this means that in this case the population of spikes on which the deviation vectors are calculated, differs between the real ZETA and random resamplings. Luckily, a simple remedy exists. We can perform “data-stitching” prior to running the random bootstrapping procedure. Specifically, between trial *k* and *k+1*, we remove all data for which

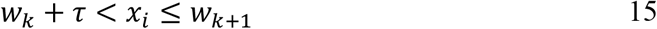

To correct for the removed time, we redefine *v* as

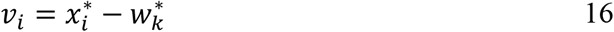

With

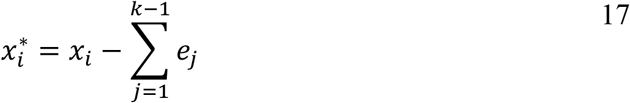

And

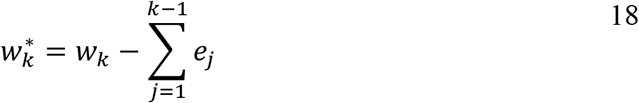

Where

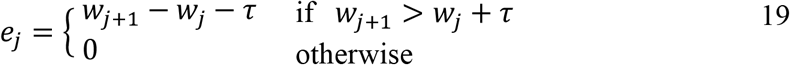

This way, jittering *w* can no longer introduce new spikes that were not also used in the calculation of *ζ*_*r*_, and our assumptions in the derivation of the ZETA-test, and specifically its time-invariance, hold (Montijn et al., 2021).

### Time-series ZETA to obtain super-sample time resolution

In contrast to the original ZETA-test, where all variables relate to point events *v*_*i*_, time-series data consist of scalar values *y*_*i*_ sampled at time points *t*_*i*_ relative to some set of events, such as stimulus onsets. Therefore, in order to adapt the ZETA-test for use on time-series data, we first need to construct a reference sample time vector with respect to stimulus onset. In many time-series data sets, the sample times are not strictly synchronized to the event times of interest; for example, one may be recording calcium imaging data at some frequency (e.g., 15.01 Hz) while the screen that presents visual stimuli updates at 59.97 Hz. This means that the delay between the sample acquisitions and the event onsets can be variable across trials. We therefore define the reference time as the set of unique sample times over all *q* trials, where for each trial *k* the sample times are relative to the most recent stimulus onset *w*_*k*_, and we set the tolerance for “uniqueness” to 1/100^th^ of the median inter-sample duration. Note that this “super-resolution” is redundant in the case of slow signals, like calcium imaging, that are recorded at a constant acquisition rate. However, using this approach, the test can also easily be applied to data with a variable acquisition rate. Moreover, it can allow a finer determination of response onsets in cases where the time-limiting factor is the acquisition rate rather than the temporal dynamics of the signal (for example voltage imaging with fast sensors).

We start the time-series ZETA-test by defining a reference time vector ***T***_*k*_ as the set of sample times *t*_*i*_ between the *k*’th event and the *k*’th event + *τ*:

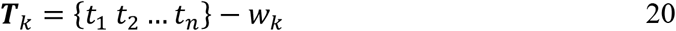

With

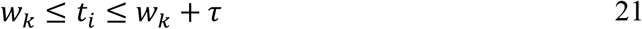

Therefore, for each event, we obtain a set of sample times, and we define the reference sample-time vector as the union over these sets:

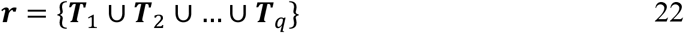

For a single trial *k* with sample times ***T***_*k*_ and values ***v***, we linearly interpolate the data to the reference time ***r***:

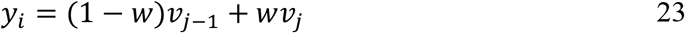

Where

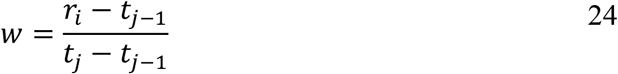

With *j* chosen such that

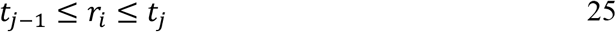

We repeat this for each of the *q* trials, obtaining a [*q* by *n*] matrix ***Y*** of data values, with *n* = |***r***| denoting the cardinality of ***r***. Taking the mean over trials to obtain 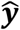 and rescaling the range to [0 1], we reduce this to a scaled vector ***u*** with *n* elements:

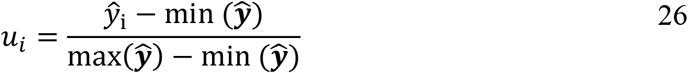

Now we replace the fractional spike position in ***g*** (Eq. 3) with a cumulative sum ***s*** over ***u***:

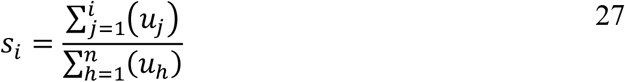

Furthermore, in the time-series case, if the values would be independent of time, the expected value at any time point is identical and equal to the average value of ***u***. Then the linear baseline density vector ***b*** (Eq. 4) corresponding to a non-modulated cumulative sum ***s***, is simply

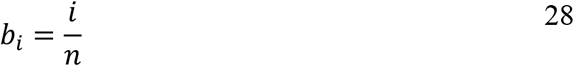

Therefore, for the time-series ZETA (T-ZETA), the temporal deviation ***δ*** is given by:

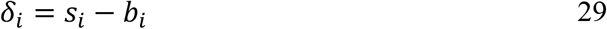

The remaining steps are identical to the original ZETA calculation (Eqs. 6-13).

### A two-sample ZETA-test

The ZETA-test can be used to detect whether the response of a neuron is modulated by sensory, optogenetic, or behavioural events. But in many cases, researchers might be interested in whether two sets of stimuli lead to a different modulation of the same neuron; or whether two neurons respond differently to the same stimulus. We therefore developed a two-sample version of the ZETA-test. In the following paragraph we will refer to condition *α* and condition *β* as two sets of data we are comparing, which can be, for example, the response of one neuron to two sets of events. The only requirement we place on the data is that the temporal window *τ* is the same for both conditions. We will therefore be working with the event times and event-relative spike times ***w***^*α*^ and ***v***^*α*^ of condition *α*, and ***w***^*β*^ and ***v***^*β*^ of condition *β*. Note that here, and in the rest of the manuscript, we use the *α* and *β* superscripts as condition indicators and not as an exponent.

A first intuitive approach to constructing a two-sample ZETA-test might be to compare the temporal deviations ***d*** (Eq. 6) between the two conditions, but this will fail when the two conditions only differ in absolute spiking rates. Since ***d*** is based on normalized spiking position ***g***, the absolute number of spikes is discarded in Equation 3. However, collapsing all spikes across trials without normalization would lead to differences in total spiking numbers when there is a difference in number of trials, but not spiking rate. We therefore construct a cumulative spiking vector for each condition based on the average number of spikes per event-window:

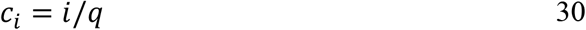

With *q* being the number of events and *n* the total number of spikes, this means that *c*_*n*_ *= n/q*, the average number of spikes per event. Note that dividing *c*_*n*_ by *τ* would therefore yield the average spiking rate per second. We perform this calculation for both conditions, obtaining ***c***^*α*^ and ***c***^*β*^. To compare these two vectors, we first create a reference time-vector ***ρ*** that contains the spike times of both ***v***^*α*^ and ***v***^*β*^:

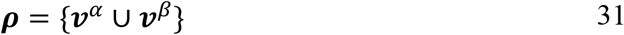

Then we linearly interpolate ***c*** to ***ρ*** for condition *α*:

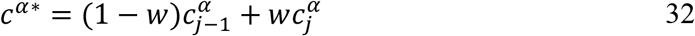

With

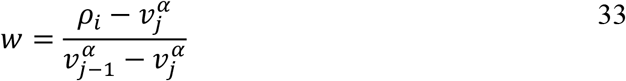

And we repeat the same procedure for c^β*^. Now we can compute the difference in cumulative spike counts by simply taking the difference between the two ***c***^*\**^ vectors:

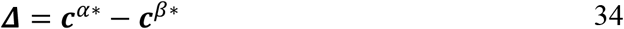

We then obtain our raw two-sample ZETA metric *Z*_*r*_:

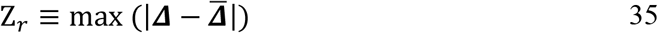

where 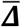 is the mean of over all spike times. Similar to the one-sample ZETA-test, we now need to normalize *Z*_*r*_ to the intrinsic variability of ***v***^*α*^ and ***v***^*β*^. Jittering works well for one-sample comparisons, but this does not satisfy the assumptions of our null hypothesis in the case of two-sample comparisons: we do not wish to know if the difference between condition *α* and *β* is larger than if they were both unmodulated. Rather, we wish to know if one modulation pattern is different from the other.

Therefore, to construct a null-hypothesis distribution of ZETA values in the two-sample test, we take random trials from the unified set of *α* and *β*. First we separate all spikes into distinct sets for each event *k*, such that

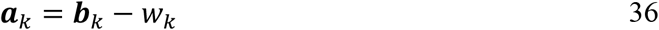

Where

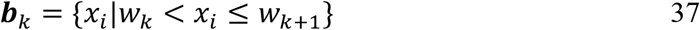

Therefore, each of the *q*^*α*^ and *q*^*β*^ events correspond to a set of event times ***a***, constituting a unified set ***A*** from which we will randomly select trials to generate our null-hypothesis samples:

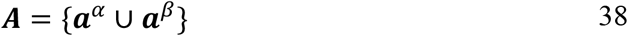

To create a random null-hypothesis sample *m* for condition *α*, we will take *q*^*α*^ random sets from ***A*** with replacement and recombine them to create a null-hypothesis version of ***v***:

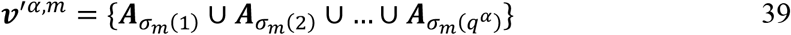

Here, ***σ***_*m*_ is a vector containing the *q*^*α*^ random integers in the range [1, *q*^*α*^ *+ q*^*β*^]. We repeat the procedure for condition *β*, and for each random sample *m*, we obtain two shuffle-randomized spike vectors ***v****’*^*α*^ and ***v****’*^*β*^. We then plug these into Equations 30 -35 to generate a null-hypothesis ZETA sample *Z’*, and use Equations 13 - 14 to transform these values into a statistical metric and corresponding p-value.

The two-sample ZETA-test does not incorporate a linear baseline subtraction like the ZETA-test, which means our original derivation of its time-invariance property no longer applies (Montijn et al., 2021). Nevertheless, it can be shown that the two-sample ZETA-test is time-invariant under the additional assumption that the difference in spiking rate between two conditions is zero at t = 0 and t = τ. These assumptions are valid in all cases where the time window and events are chosen appropriately. Whenever a time-locked difference exists at t = 0, this means the events of interest cause a change in a cell’s response before the onset of the stimulus, and the event times are either incorrect, or the experiment is performed in a time machine. On the other hand, whenever a time-locked difference exists at t = τ, this means that the window length was too short, and τ should be increased. In other words: if the underlying data are sound, the two-sample ZETA-test is time-invariant.

### The one-sample ZETA-test is a special case of the two-sample ZETA-test

One can find an equivalence between the one- and two-sample tests in the case where condition *α* is a set of events, and condition *β* represents the linear baseline. This linear baseline can be approximated by an infinite set of events occurring over the same time window as the events in condition *α*. This means that all temporal modulation is averaged out, and the resulting cumulative spike count of condition *β* becomes identical to linear baseline density vector ***b*** from Equation 4 if sampled at ***v***^*α*^, and divided by *τ*. Randomly selecting *q*^*α*^ sets from ***A*** (Eq. 38), when *q*^*β*^ = ∞, therefore means selecting *q*^*α*^ random event-times, which is identical to jittering event onsets with an infinite jittering window. For condition *β* we take infinite samples from infinite events, so ***v***^*β*^ will remain identical to linear baseline density vector ***b***. Subtracting these vectors yields ***δ****’* (Eq. 11). The steps thereafter (Eq. 12-14) are the same for the one-sample and two-sample tests, so they are therefore equivalent under these conditions.

Note that sampling at ***v***^*α*^ is only strictly necessary to make the one-sample test computationally tractable, but this does not influence its mathematical behaviour. Moreover, a division by *τ* can be added to the two-sample test without affecting its statistical properties – it merely scales the values in ***A***. The only real difference between the one-sample-equivalent version of the two-sample test, and the actual one-sample test is therefore the width of the jittering window. In data sets that are non-stationary across event repetitions, the local jittering of the one-sample test ensures uniform sampling over the signal’s non-stationarity, but this is absent from the two-sample zeta-test. On the other hand, when signals are stationary across events, the one-sample-equivalent two-sample test and the one-sample test are truly mathematically equivalent.

### Two-sample time-series ZETA-test

Having described the time-series ZETA-test and the two-sample ZETA-test, the two-sample time-series ZETA-test is a straightforward combination of these procedures. Rather than calculating the deviation vector ***δ*** between the cumulative sum ***s*** and linear baseline vector ***b***, we calculate a cumulative sum for both condition *α* and condition *β*. We replace Eqs. 24-26 with

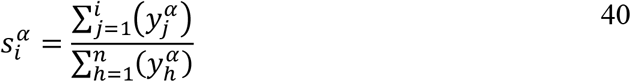

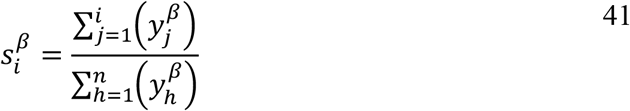

Then the two-sample deviation vector ***Δ*** becomes:

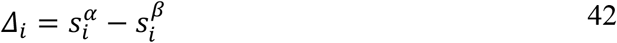

Now our raw two-sample time-series ZETA metric *Z*_*r*_ can be calculated as before by taking the maximum of the absolute of the mean-normalized deviations. To generate null-hypothesis samples, we repeat the above steps, but construct a [*q* by *n*] matrix ***Y*** for both conditions by randomly selecting *q*^*α*^ or *q*^*β*^ trials from the unified set of *q*^*α*^ + *q*^*β*^ possible trials for each random sample. Finally, we obtain the statistical metric using Eq. 13 - 14.

### GCaMP6f and Neuropixels data

The calcium imaging data analysed in this paper are the same as previously described (Montijn et al., 2016). We used GCaMP6f data from 15 recordings in four C57BL/6 mice. Cell bodies were detected semi-automatically using an open-source toolbox (https://github.com/JorritMontijn/Preprocessing_Toolbox) and cells were only included for further analysis if their somata were clearly distinguishable from the background neuropil. The electrophysiological data used here are also previously described elsewhere (Montijn et al., 2023). In short, we performed 21 repeated-insertion recordings in seven C57BL/6 mice with Neuropixels. All mice were housed in a 12 h/12 h dark/light cycle with ad libitum access to food and water and were awake during recording. The recording setup presented drifting gratings of 24 directions (spaced in 15-degree steps) and was controlled using Acquipix (Montijn, 2022). Spikes were sorted post-hoc using Kilosort 2.5 (Pachitariu et al., 2023), and electrode location was determined by aligning histological slices and neurophysiological landmarks to the AllenCCF mouse brain atlas (https://github.com/cortex-lab/allenCCF) using the UniversalProbeFinder (Montijn & Heimel, 2022). All code used in the Neuropixels data acquisition and pre-processing is available online in the Acquipix and UniversalProbeFinder repositories on https://github.com/JorritMontijn. We included only clusters of sufficient quality, as quantified by their spike contamination ratio and Kilosort’s quality labelling, for further analysis. For more detailed information, see (Montijn et al., 2023). All experiments were approved by the animal ethics committee of the University of Amsterdam (GCaMP6f) and Royal Netherlands Academy of Arts and Sciences (Neuropixels), in compliance with all relevant ethical regulations.

### ECoG data

The example application to ECoG data was performed on a freely available dataset (Miller et al., 2016). All patients participated in a purely voluntary manner, after providing informed written consent, under experimental protocols approved by the Institutional Review Board of the University of Washington (#12193). All patient data were anonymized according to IRB protocol, in accordance with HIPAA mandate.

### Experimental benchmarks

We verified the statistical performance of the one-sample ZETA test and one-sample time-series ZETA test tests by calculating the p-value for stimulus responsiveness for all neurons. The false positive rate was computed by repeating the procedure after jittering the stimulus onset times. For the two-sample comparisons, we calculated whether each neuron was differentially responsive to the presentation of two directions of a drifting grating, or to two different scenes in a natural movie. We calculated the false positive rate by randomly selecting 50% of spikes from either condition to create new, randomized conditions. For the time-series-based two-sample tests, we used a similar procedure, except we took 50% of trials rather than spikes to compute the false-positive rate (as selecting a random 50% of samples would change temporal autocorrelations).

### Synthetic benchmarks one-sample ZETA-test

We quantified the effect of the data-stitching procedure by generating spiking responses of two types of synthetic neurons with exponential inter-spike intervals as firing distributions. For the first type of neuron (fig. 1G), we generated 160 trials that varied in duration, ranging from 0.5 to 10 seconds, and partially overlapped. Each trial consisted of an onset period of 100 ms, followed by a 900 ms sustained period, after which the neuron returned to its baseline rate. For each neuron the baseline rate was chosen to be λbase = 0.1 + Exp(0.1), the onset rate λonset = 0.2 + Exp(1), and the sustained rate λsust = Exp(0.1), where Exp(*k*) is a random variable drawn from an exponential distribution with parameter *k*. We simulated a second type of neuron to illustrate the effect that stitching versus no-stitching can have (fig. 1H). Again, we used 160 trials, all of 1 second in duration, with a 3 second inter-trial period. The base rate of this neuron during inter-trial intervals was Exp(0.1)+0.1, the onset rate (100 ms) was Exp(4)+4, and the sustained rate (900 ms) was Exp(2)+2. Note that by setting the ZETA-test’s trial duration τ to 1 second, ZETA’s real deviation does not cover the transition into and out of the baseline firing rate, while the jittered deviations will include one of these transitions.

### Synthetic benchmarks two-sample tests

To further investigate the statistical properties of our methods, we also used two types of generated synthetic data sets. In both cases, we generated neurons with a baseline rate as described above; an exponential inter-spike interval model with a rate randomly chosen from λ∼ Exp(1). Furthermore, one additional spike was added to half of all trials on top of these baseline spikes. The timing of these spikes followed a Gaussian distribution with a mean of 55 ms and a standard deviation of 1 ms. This therefore simulates a cell with a sharp onset peak.

For the first benchmark, we generated a neuron with the same baseline firing rate, but which had an onset spike in all trials – it therefore differed in both peak height and total number of spikes (owing to the higher peak). For the second benchmark, we instead generated a neuron with the same baseline firing rate and same amount of onset spikes, but whose onset peak was instead shifted by 2 ms – this means the neurons differed in onset peak latency but not in total number of spikes. In both cases, the two-sample tests (ZETA, ANOVA, t-test) were then applied to obtain a p-value. False positive rates were calculated by generating the second neuron with the exact same parameters as the first neuron: the same number of onset spikes occurring at the same peak time.

### ANOVAs and (optimal) bin width selection

We determined the optimal bin size for the ANOVA using the Shimazaki & Shinomoto procedure, which calculates a loss value for a set of trial-locked spike times and a chosen bin width (Shimazaki & Shinomoto, 2007). The binning width was optimized separately for each individual neuron using a loss minimization function. We determined the bin width for two-sample comparisons by combining the spike times from both conditions. ANOVAs performed on time-series data used the samples of the time-series data themselves, so no bin width optimization was necessary. The calcium imaging data sets had a sample duration of 79 ms (12.7 Hz). The ECoG data had a sample duration of 1 ms. One-sample comparisons were performed using a one-way ANOVA across bins, with the reported p-value being the single main effect. Two-sample comparisons were performed using a two-way ANOVA on bins x condition. The reported p-value was the lower of the two Bonferroni-corrected p-values for *condition* main effect and *bin x condition* interaction.

### The Maris & Oostenveld clustering test

The Maris & Oostenveld clustering procedure originally produces a sum of t-statistics for each cluster of bins that exceeds a statistical threshold. This threshold is a hyperparameter that needs to be chosen by the experimenter, but a common approach is to set this at α = 0.05 (Maris & Oostenveld, 2007). Each cluster’s sum of t-statistics is then compared to a null-hypothesis distribution obtained by randomly combining trials from the two conditions and repeating the clustering procedure. Each cluster therefore receives its own p-value, so to obtain a single estimate for any difference between two conditions, we discarded all but the cluster with the lowest p-value. This procedure is strictly one-tailed, so to be able to detect both positive and negative peaks, we ran the procedure twice and took the lowest of the Bonferroni-corrected positive-peak and negative-peak p-values (fig. 5N). Therefore, as we used 1000 random iterations, the lowest possible p-value of this “Clust-t” test was p = 0.002. Open-source code for this test can be found here: https://github.com/JorritMontijn/GeneralAnalysis/blob/master/statistics/clustertest.m.

### Inclusion error metric

We calculated the excess in false positive rate (FPR) of all ZETA-tests, t-tests, and ANOVAs applied to experimental data with the aim to quantify the inclusion errors we observed in ANOVA’s behaviour. This metric *f* is a log-normalized integral over the excess FPR relative to the theoretical norm (FPR/α = 1), and takes values in the range [0,]. It scales such that *f* = 0 for FPR/α ≤ 1, while *f* = 1 means that FPR/α = *e* (Euler’s number, ≈ 2.7). The metric acts on a sorted set of *n* randomized control p-values ***v***:

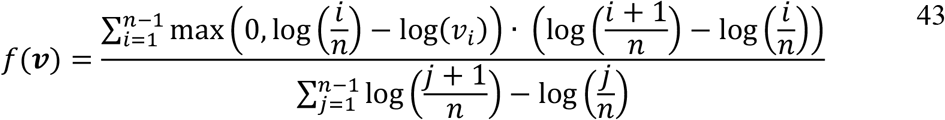

Note that we constructed the metric to work in logarithmic space to more accurately reflect the logarithmic plots of FPRs we present throughout the manuscript. We chose logarithmic space in the analysis and presentation to emphasize smaller p and α values, as the difference between p = 0.001 and p = 0.101 is often of more importance than the difference between p = 0.701 and p = 0.801.

## Code Accessibility

URLs for specific analyses are stated throughout the manuscript. In general, all code is open-source and can be accessed here: https://github.com/JorritMontijn. The MATLAB implementation of the ZETA-tests can be found here: https://github.com/JorritMontijn/zetatest. The Python implementation can be downloaded using PyPi or from GitHub: https://github.com/JorritMontijn/zetapy.

## Acknowledgements

We thank Camilo Rojas, Valeria Gazolla and Christian Keysers for suggesting to pursue a ZETA-test for time-series data, and Matt Self for providing an initial implementation of the Maris & Oostenveld clustering procedure. We would also like to thank the following people for their feedback on GitHub: Rebecca Mease, Jonah Pearl, Magdalena Sabat, Rejwan Salih, Marina Slashcheva, and Wolf De Wulf. This work was funded by a Royal Netherlands Academy of Arts and Sciences (KNAW) Fonds KNAW-Instituten grant.

